# Intratumoral Heterogeneity Promotes Collective Cancer Invasion Through NOTCH1 Variation

**DOI:** 10.1101/2021.06.30.450540

**Authors:** Peter Torab, Yue Yan, Mona Ahmed, Hironobu Yamashita, Joshua I. Warrick, Jay D. Raman, David J. DeGraff, Pak Kin Wong

## Abstract

Cellular and molecular heterogeneity within tumors has long been associated with the progression of cancer to an aggressive phenotype and a poor prognosis. However, how such intratumoral heterogeneity contributes to the invasiveness of cancer is largely unknown. Here, using a multidisciplinary approach, we investigate the interaction between molecular subtypes within bladder microtumors and the corresponding effects on their invasiveness. Our results reveal heterogeneous microtumors formed by multiple molecular subtypes possess enhanced invasiveness compared to individual cells, even when both cells are not invasive individually. To examine the molecular mechanism of intratumoral heterogeneity mediated invasiveness, live single cell biosensing, RNA interference, and CRISPR-Cas9 gene editing approaches were applied to investigate and control the composition of the microtumors. An agent-based computational model was also developed to evaluate the influence of NOTCH1 variation on DLL4 expression within a microtumor. The data indicate that variation in NOTCH1 expression can lead to upregulation of DLL4 expression within the microtumor and enhancement of microtumor invasiveness. Overall, our results reveal a novel mechanism of heterogeneity mediated invasiveness through intratumoral variation of gene expression.

**Summary statement:** This study reveals a mechanism that Notch1 variation, instead of the average value, promotes the invasiveness of microtumor, providing a link between intratumoral heterogeneity and collective cancer invasion.

## Introduction

In the United States, bladder cancer is a common malignancy, with an estimated 83,730 people diagnosed and over 17,200 individuals dying annually (Siegel et al., 2021). At both the cellular and molecular levels, bladder cancer is an extraordinarily heterogeneous disease. At the individual patient level, several groups have independently completed molecular characterization of human bladder cancer (Cancer Genome Atlas Research, 2014; Choi et al., 2014; Damrauer et al., 2014). These efforts led to the discovery of luminal, basal/squamous, and other transcriptional subtypes (stroma-rich and neuroendocrine-like) of the muscle-invasive bladder cancer (Kamoun et al., 2020). Luminal bladder cancer (so called because of expression of luminal urothelial markers) is often associated with histology consistent with urothelial cell carcinoma and is further classified into luminal papillary, luminal nonspecified, and luminal unstable (luminal papillary is the most common molecular subtype). Luminal cancer cells are controlled by the transcriptional master regulators, such as FOXA1, PPARγ, and GATA3 (Warrick et al., 2016; Yamashita et al., 2019). Luminal cancer cells are reportedly chemoresistant (Choi et al., 2014), while a subset of these tumors appear to respond favorably to immune checkpoint blockade (Mariathasan et al., 2018). On the other hand, the basal/squamous subtype of bladder cancer is enriched for morphologic squamous differentiation. Such basal/squamous bladder cancers are highly lethal (Choi et al., 2014). Basal/squamous cancer cells are typically classified by elevated expression of specific cytokeratins, such as KRT5 and KRT14 (Warrick et al., 2016). While it appears that basal/squamous tumors respond favorably to neoadjuvant, platinum-based chemotherapy and targeted therapeutic approaches, recent reports indicate patients with basal/squamous bladder cancer have inferior overall and disease-specific survival rates (Choi et al., 2014).

While associations between transcriptional subtypes and clinical outcome have been observed at the patient level, the impact of intratumoral heterogeneity is similarly associated with aggressive disease characteristics. For example, tumors with intratumoral cellular heterogeneity usually present at advanced stage and exhibit poor clinical outcome (McGranahan and Swanton, 2015). Relative to pure urothelial cell carcinoma, urothelial carcinoma with squamous differentiation often presents as high-stage disease with lymph node metastasis (Liu et al., 2017; Warrick et al., 2019). It has been suggested that distinct cell subpopulations may cooperate as a community to support cancer progression. For example, cancer cells with invasive characteristics can promote the invasion of non-invasive epithelial cells in a co-culture spheroid model of breast and prostate cancer (Carey et al., 2013). Basal cancer cells can serve as invasive leader cells to promote collective invasion (Cheung et al., 2013; Hwang et al., 2019). Furthermore, NOTCH1, which suppresses basal phenotypes and bladder cancer progression (Goriki et al., 2018; Maraver et al., 2015; Rampias et al., 2014), negatively regulates the formation of a DLL4 expressing subpopulation that promotes collective cancer invasion (Dean et al., 2016; Riahi et al., 2015). In a 3D invasion model of bladder cancer, microtumors with a high level of DLL4 at the invasive front exhibited enhanced invasiveness (Torab et al., 2020). Overall, the available evidence suggests non-cell autonomous interactions between heterogeneous cell populations can enhance the aggressiveness of cancer and underscores heterogeneous cell populations may collectively promote cancer invasion. Nevertheless, the influence of intratumoral heterogeneity on bladder cancer cell behavior remains poorly understood, limiting innovation in the clinical management of this common disease.

In this study, we investigate the influence of tumor heterogeneity on the invasiveness of cancer by establishing a tumor bioengineering approach (**Fig. 1a-b**). The tumor bioengineering approach simultaneously generates a large number of microtumors in a single assay and recapitulates the important features of the tumor microenvironment, including multilayer extracellular matrix (ECM) components and heterogeneous molecular subtypes of cancer cells. In addition, we incorporate a locked nucleic acid (LNA) single cell biosensor to perform *in situ* gene expression analysis in 3D microtumors. Using established cell lines with luminal and basal/squamous signatures, transient knockdown by siRNA, CRISPR knockout, and *in silico* models, we examine the influence of heterogeneity of NOTCH1-DLL4 signaling on bladder cancer invasion. The results suggested that variation, or molecular heterogeneity, of NOTCH1 expression within the microtumor can promote the invasiveness of heterogeneous bladder cancer. Overall, our tumor bioengineering approach reveals a novel mechanism of heterogeneity mediated invasiveness in bladder cancer.

**Figure 1.**
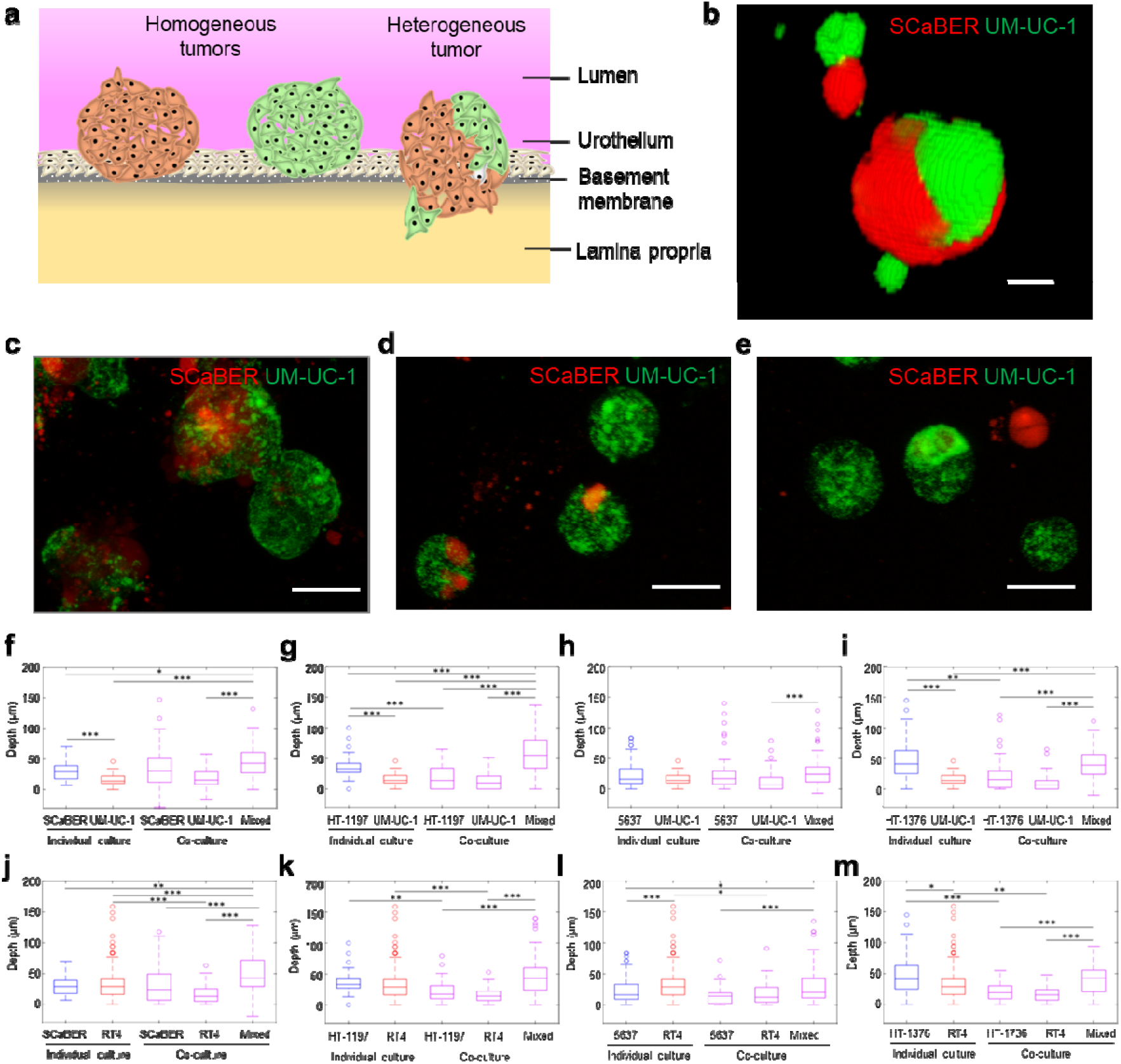
Co-culture microtumors invading extracellular matrix model. (a) Schematic of heterogeneous microtumor invasion through basement membrane and lamina propria. (b) 3D reconstruction of heterogeneous microtumor created using NIH ImageJ 3D viewer plugin. Scale bar, 10 μm. (c-e) Vertical projection views of SCaBER-UM-UC-1 heterogeneous microtumors. Scale bars, 50 μm. (f-m) Invasion depth into basement membrane ECM for basal- (blue) and luminal-type (red) microtumors, as well as co-culture (purple) microtumors. Co-culture results are separated for homogeneous and heterogeneous (mixed) microtumors within co-culture experiments. *p<0.05, **p<0.01, ***p<0.001 Kruskal-Wallis ANOVA test with Tukey-Kramer post-hoc test.

## Results

### Cellular heterogeneity enhances invasiveness of bladder microtumors

The presence of multiple histologic variants within a bladder cancer tumor is common, and molecular heterogeneity is often observed when multiple variants are present (Warrick et al., 2019). To study these effects, cell lines representing two most common bladder cancer subtypes, i.e., luminal papillary and basal/squamous, were co-cultured to form mixed (i.e., heterogeneous) microtumors (Kamoun et al., 2020). The single layer invasion model consisting of simulated basal membrane matrix was first applied to test cell lines representing the luminal papillary subtype (UM-UC-1, RT4) and basal/squamous subtype (SCaBER, HT-1197, HT-1376, 5637) individually and in every possible basal-luminal combination. The cells were self-assembled on the ECM mimicking gel and formed 3D microtumors approximately 50-100 μm in diameter (**Supplementary Movies 1-3**). For co-culture experiments, cell lines were stained separately, then mixed at a 1:1 ratio (**Fig. 1b**). Within mixed microtumors, individual cell types tended to aggregate together (**Fig. 1c**). Microtumors composed of both cell types (mixed or heterogeneous) and a single cell type (homogeneous) were observed in the co-culture experiment (**Fig. 1d-e**).

The 3D invasion assay allowed microtumors to invade into the ECM-mimicking gel. In accordance with our previous report (Torab et al., 2020), the invasion depth correlated the invasiveness of the cancer cell lines determined by animal models and other 3D invasion assays (**Fig. 1f-m**) (Fujiyama et al., 2001; Jager et al., 2014; Luo et al., 2016; Zuiverloon et al., 2018). For example, non-invasive cell lines, such as UM-UC-1 (derived from grade 2 bladder cancer and classified as the luminal papillary subtype based on the consensus molecular classification of muscle-invasive bladder cancer) (Jager et al., 2014; Kamoun et al., 2020), produced microtumors that displayed only slight deformation of the matrix. The depth of deformation is typically less 50 μm and can be understood by contact mechanics (Style et al., 2013). In contrast, invasive cell lines, such as HT-1376 (derived from grade 3 bladder cancer with a basal/squamous signature based on the consensus molecular classification of muscle-invasive bladder cancer), displayed a relatively high invasiveness as indicated by the invasion depth (Kamoun et al., 2020; Ramakrishnan et al., 2018).

Heterogeneity in invasiveness among microtumors has been reported in organoid invasion models (Padmanaban et al., 2020). Similarly, we observed only a portion of microtumors penetrated the gel despite the microtumors were formed by a single cell line. We, therefore, characterized the fraction of microtumors which invaded into the gel. In this study, the invasive fraction is defined by the portion of microtumors with an invasion depth over 50 μm. This threshold value is consistent with our previous study (Torab et al., 2020), and a similar threshold value was also obtained by clustering analysis (**Supplementary Fig. S1**). The invasive fraction showed a similar trend compared to the invasive depth (**Supplementary Fig. S1**). In particular, UM-UC-1 had no (0%) invasive fraction while HT-1376 had a high fraction (~30%) of invasive microtumors. Notably, most cell lines exhibited a small fraction (5-10%) of invasive microtumors despite having a low level of median invasion depth, suggesting heterogeneity within individual cell lines.

We examined the influence of combining luminal-basal cells on the invasiveness of microtumors (**Fig. 1f-m** and **Supplementary Fig. S2**). Remarkably, we observed an increase in invasiveness in microtumors with mixed luminal and basal cell types compared to homogenous microtumors formed by individual cell lines. The median invasion depth of mixed microtumors was larger than most homogeneous microtumors. There was a significant increase (one-tailed Wilcoxon rank sum test) in the median invasion depth of mixed microtumors compared to the majority of basal cell lines (75%) and luminal cell lines (88%) that comprised the mixed microtumors. For instance, both the invasion depth and the invasive fraction of mixed SCABER/UM-UC-1 microtumors were significantly enhanced compared to SCaBER and UM-UC-1 cells cultured individually (**Supplementary Fig. S3**). For invasive cells, such as HT-1376, similar or slightly higher invasion depths and invasive fraction were observed when mixed with UM-UC-1 or RT4, which were less invasive individually. We also analyzed the influence of the microtumor composition on the invasiveness as both homogeneous and heterogeneous microtumors were formed in the co-culture experiments. Heterogeneous microtumors showed a larger invasion depth compared to most homogeneous microtumors in co-culture experiments (**Fig. 1f-m**). Since both basal and luminal cells were in the same well, the enhancement in invasiveness could not be explained fully by diffusible factors and was likely contributed by a contact dependent mechanism. Taken together, these results support the notion that the presence of heterogeneous subtypes enhances bladder cancer invasion via a contact dependent mechanism. The combination of SCaBER and UM-UC-1 cells was selected as the model for subsequent experiments, as both the invasion depth and invasive fraction were enhanced considerably compared to the individual cell lines.

### Heterogeneous microtumors efficiently invade matrices found in the bladder wall

We further evaluated the behaviors of mixed microtumors by establishing a multilayer invasion model. The materials and gelation procedures were optimized to create a microenvironment that mimics the basement membrane and lamina propria. Specifically, gelation of a thick layer of collagen I was first allowed for simulating the lamina propria. The mechanical properties and porosity of the collagen I layer was optimized for probing the invasiveness of cancer (Nam et al., 2016; Tien et al., 2020; Wisdom et al., 2018; Zaman et al., 2006). After gelation, a thin layer of Matrigel was loaded on top of the collagen similar to the single layer invasion model. The Matrigel layer is thicker compared to the bladder basement membrane for uniform coverage and visualization of the microtumor invasiveness. The multilayer model also allowed the generation of a gradient of serum or other chemicals for promoting directional invasion of microtumors. In this study, an FBS gradient was created across the layers to simulate a nutrient gradient in the bladder wall. Similar to the single layer invasion model, microtumors were self-assembled and allowed to invade into the matrices (**Fig. 2a**). The use of the multilayer invasion model can reveal the microtumor’s ability to breach the ECM protein found in the basement membrane and the underlying lamina propria (**Fig. 2b**). The ability of cells to progress through both of these layers would indicate that the tumor has a higher chance to progress into muscle invasive bladder cancer, while non-muscle invasive tumors would not be expected to penetrate through the ECM mimicking gel.

**Figure 2.**
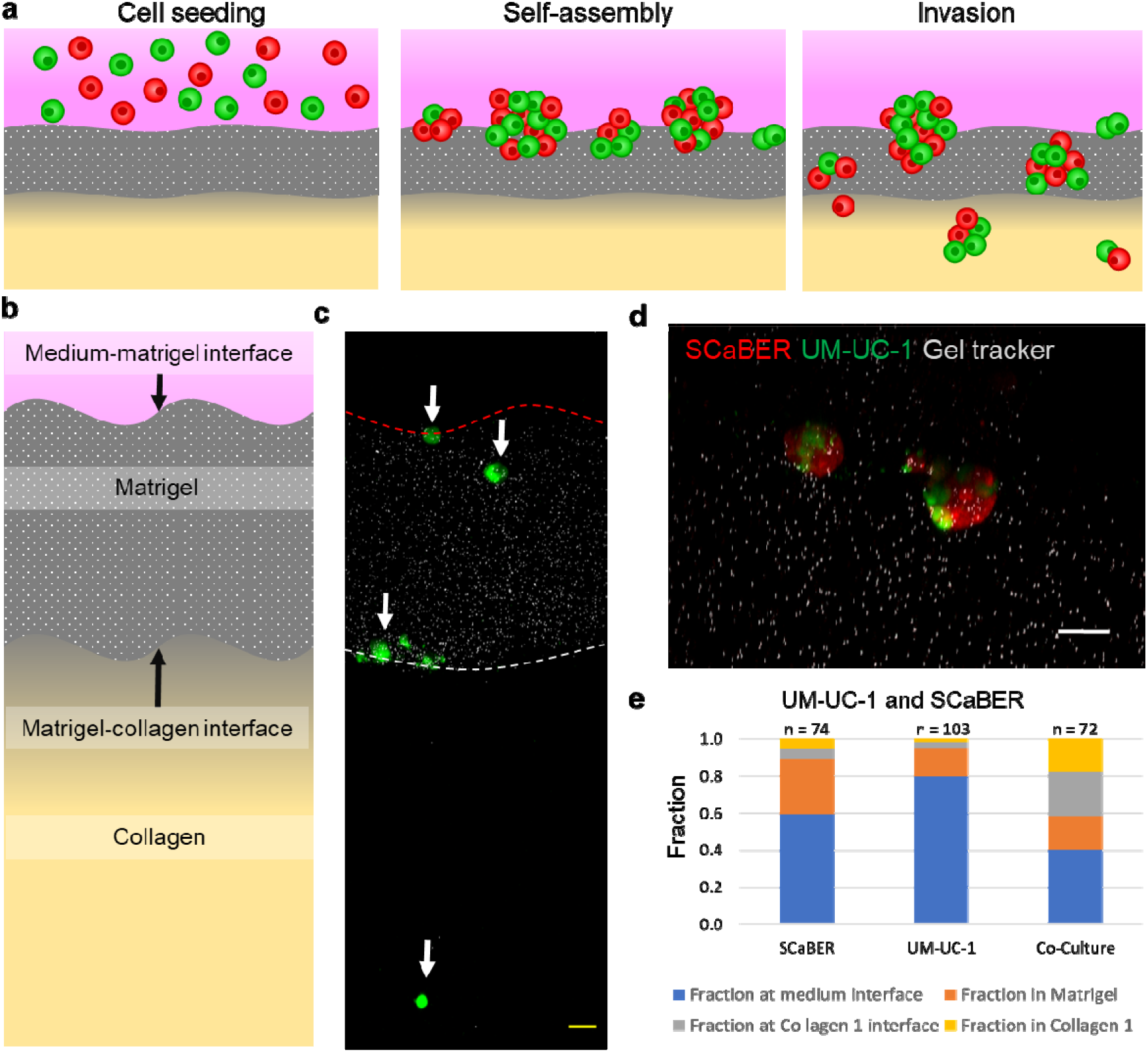
Heterogeneous microtumors invading multilayer ECM. (a) Schematics of self-assembly and invasion of heterogeneous microtumors. (b) Schematic of multilayer ECM model. (c) Cross-section view of multilayer ECM. White arrows indicate microtumors in each of the four regions. Red dotted line indicates the medium-Matrigel interact, and white dotted line indicates the Matrigel-collagen interface. (d) Heterogeneous microtumor invading the Matrigel layer. Scale bars, 50 μm. (e) Fraction of microtumors counted in each region on the multilayer invasion model. Results represent the total from 6 independent experiments.

In the experiment, the cells formed microtumors in a manner similar to the single layer invasion model and invaded into different regions of the gel layers, becoming embedded at the interfaces between layers or within the gel layers (**Fig. 2c-d**). For homogeneous culture, SCaBER and UM-UC-1 did not display a high invasiveness, and the majority of microtumors stayed at the medium-Matrigel interface or embedded within the Matrigel. Only a very small fraction (less than 5%) of microtumors were able to reach the collagen layer (**Fig. 2e**). In contrast, co-culture microtumors exhibited an enhanced capability of invading into the matrices. Some mixed microtumors (~20%) penetrated through the Matrigel layer and reached the Matrigel-collagen interface. Furthermore, a substantial fraction (~20%) reached into the collagen layer, suggesting an ability to invade through both Matrigel and collagen. These observations further support the notion that heterogeneous microtumors possess enhanced invasiveness.

### NOTCH1-DLL4 signaling in invasive microtumors

We investigated the mechanism that drives the enhanced invasiveness of mixed microtumors. In particular, we studied the role of NOTCH signaling, which mediates contact-dependent signaling between cells, on bladder cancer invasiveness. Loss of NOTCH1 is associated with the basal subtype (Maraver et al., 2015; Rampias et al., 2014), and NOTCH1-DLL4 signaling is shown to regulate collective cancer invasion (Dean et al., 2016; Riahi et al., 2015; Torab et al., 2020). We first evaluated the expressions of NOTCH1 and DLL4 in UM-UC-1 and SCaBER microtumors using a live single cell biosensor, which was demonstrated in cancer cells, tumor organoids derived from patients, and tumor tissues (Tao et al., 2014; Torab et al., 2020; Wang et al., 2015). In particular, locked nucleic acid (LNA) sequences targeting NOTCH1 and DLL4 mRNA were mixed with gold nanorods (GNR) and applied to SCaBER and UM-UC-1 cells (**Supplementary Fig. S4**). The cells were then seeded into the multilayer invasion model and imaged after 72 hours to measure the target mRNA expression (**Fig. 3a**). **Fig. 3b-c** show the average intensities of SCaBER, UM-UC-1 and mixed microtumors. UM-UC-1 expressed a high level of NOTCH1 mRNA relative to SCaBER, which was barely detectable. In mixed microtumors, the average NOTCH1 expression showed an intermediate value between the expression values for individual cell lines. Notably, a large variation of NOTCH1 mRNA expression was observed within the microtumor. Some cells in the microtumor displayed a high level of NOTCH1 expression while some cells had no detectable NOTCH1 expression (**Fig. 3a**). This observation was not surprising as UM-UC-1 and SCaBER expressed high and low levels of NOTCH1, respectively. In contrast, DLL4 expression in co-culture was higher than either individual cell line. This observation is interesting as DLL4 expression in mixed microtumors was not an ensemble average of the two cell lines and implicates non-autonomous regulation of DLL4 expression in mixed microtumors.

**Figure 3.**
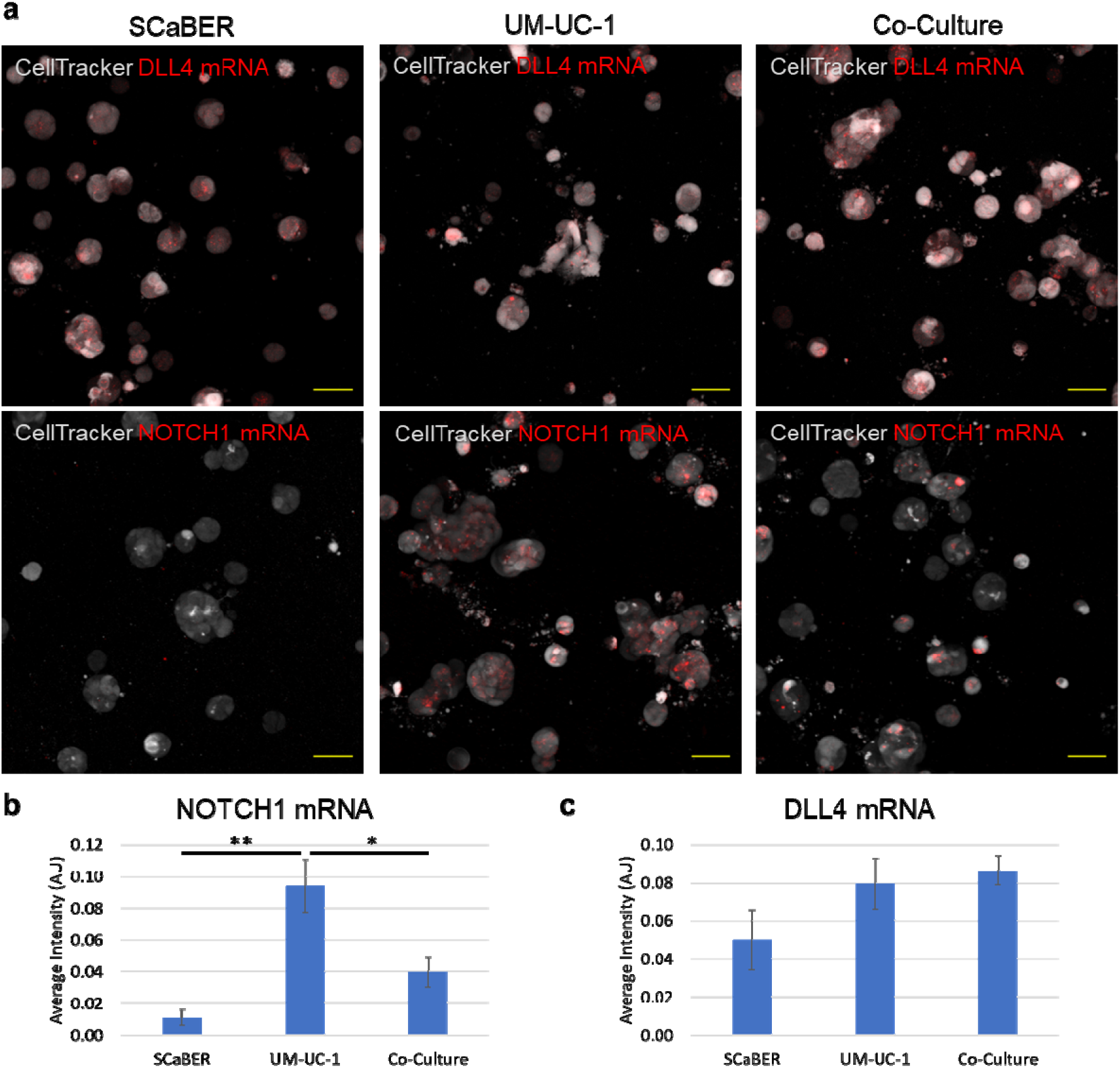
Detection of NOTCH1 and DLL4 mRNA expressions and variations using the GNR-LNA biosensor. (a) Representative images of biosensors in SCaBER, UM-UC-1, and co-culture microtumors. (b-c) Fluorescence intensity of (b) NOTCH1 and (c) DLL4 biosensors. *p<0.05, **p<0.01, One-way ANOVA test with Tukey’s post-hoc test. Results are representative of three independent experiments.

To test the whether the enhanced invasiveness of mixed tumors is associated with NOTCH1-DLL4 signaling, NOTCH1 and DLL4 siRNA were applied in either UM-UC-1, SCaBER or both cell types (**Fig. 4a-b**). Cells were each alternately treated with NOTCH1, DLL4, and non-targeting (control) siRNA before self-assembly. In the experiment, siRNA inhibition of DLL4 in individual cell lines and in both cell lines in co-culture substantially inhibited the cells’ ability to invade through the Matrigel layer (**Fig. 4c**). DLL4 siRNA reduced invasion when applied to either cell line or both cell lines with similar efficiency. Most microtumors stayed at the medium-Matrigel interface or in the Matrigel, and no cells were able to reach into the collagen layer (i.e., 0%). This observation is consistent with the view that DLL4 supports collective invasion of cancer (Dean et al., 2016; Riahi et al., 2015; Torab et al., 2020). On the other hand, NOTCH1 transient knockdown was performed on individual cell line or both cell lines in the co-culture invasion assay (**Fig. 4d**). When applied to either cell line, NOTCH1 knockdown resulted in an increase in invasiveness. The fraction of microtumor reaching the collagen layer increased for NOTCH knockdown in either cell line (~30%). No microtumors were observed to stay at the Matrigel-collagen interface. The fraction of microtumors in collagen was higher than co-culture without transient knockdown (~20%). Interestingly, inhibition of NOTCH1 in both UM-UC-1 and SCaBER reduced the fraction invaded into collagen (~5%), which essentially eliminated the co-culture-mediate invasiveness. Some of the microtumors were trapped at the Matrigel-collagen interface. Together, NOTCH1-DLL4 signaling is associated with the ability of microtumor to penetrate through collagen, and the enhanced invasiveness of mixed tumors correlated with the variation of NOTCH1 expression within the microtumor.

**Figure 4.**
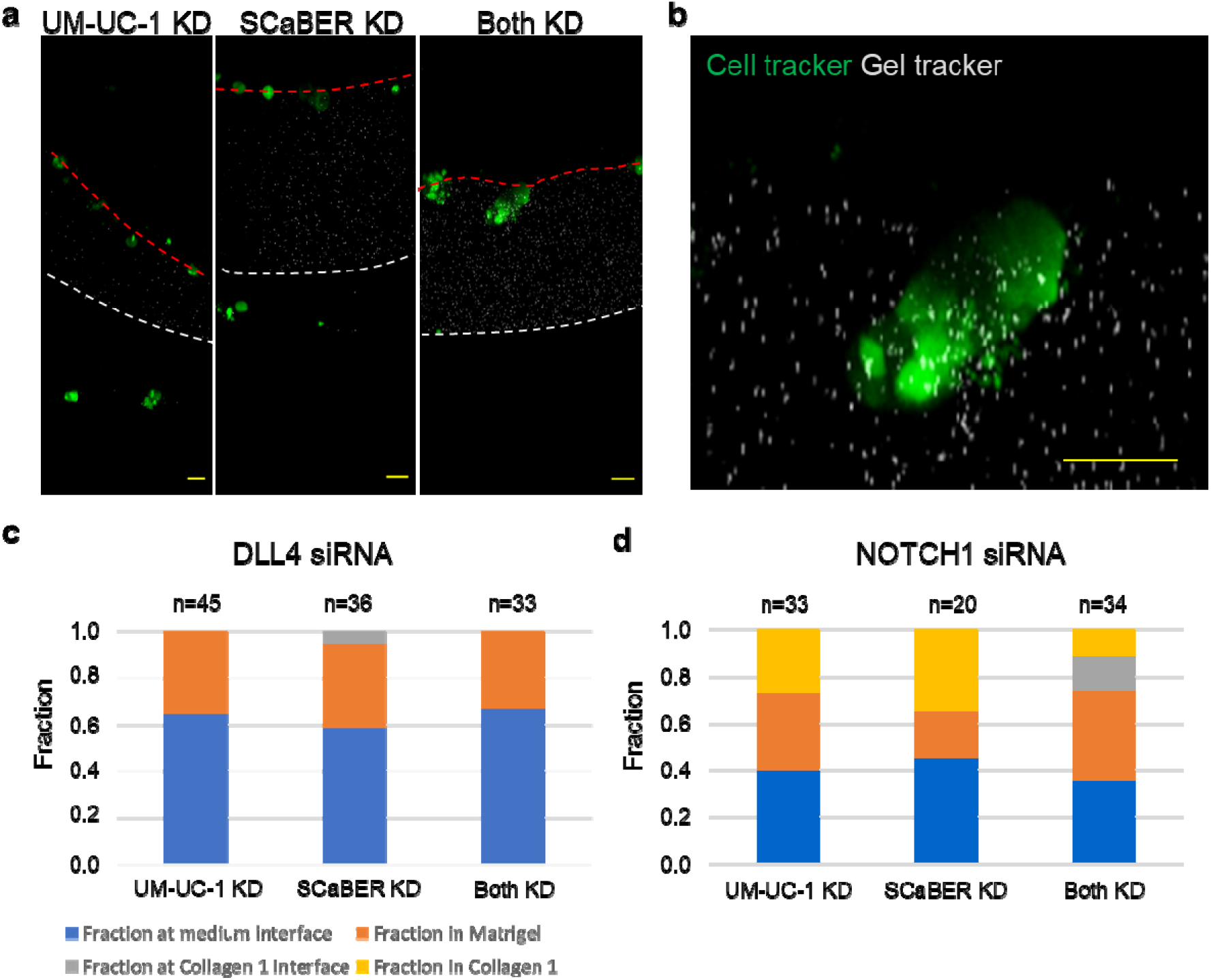
RNA interference applied to DLL4 and NOTCH1. (a) Representative images of microtumors treated with NOTCH1 siRNA. (b) Detail view of invasive microtumor. (c-d) Fraction of microtumors counted in each region with (c) DLL4 siRNA and (d) NOTHC1 siRNA. Results represent the totals from three independent experiments.

### FOXA1 knockout promotes invasion of mixed microtumors

The influence of luminal-basal co-culture on microtumor invasiveness was further investigated by CRISPR-Cas9 gene editing. FOXA1 is an emerging regulator of the luminal subtype of bladder cancer, and FOXA1 overexpression can drive basal bladder cancer cells to assume a more luminal phenotype (Warrick et al., 2017; Warrick et al., 2016). FOXA1 has also been implicated in the regulation of NOTCH1 in other cancer types (Qiu et al., 2014). A FOXA1 knockout (KO) cell line was generated from the luminal UM-UC-1 cell line using CRISPR-Cas9 gene editing. The FOXA1-KO UM-UC-1 cell line exhibited a reduced expression of NOTCH1 and NOTCH1 targeted genes, such as HEY1, HES1, and CDKN1A (p21) compared to the wildtype cell (**Fig. 5a**). The invasiveness of the UM-UC-1 FOXA1 KO cells was tested in the multilayer invasion model. The KO cells showed a slight increase in the fraction of microtumor reaching the collagen interface while the overall invasive fraction was similar between the wild-type and FOXA-KO UM-UC-1 cells (**Fig. 5b**). UM-UC-1 FOXA1-KO and UM-UC-1 wild type cells were co-cultured to form microtumors in the multilayer invasion model (**Fig. 5b**). Intriguingly, mixed microtumors were much more invasive than microtumors formed by wild type or KO cells individually. Both the overall invasive fraction and the fraction of microtumor invaded into collagen increased by co-culture of the cells. Furthermore, the mixed microtumors were less likely to be trapped at the Matrigel-collagen interface. These results further support the notion that variation of NOTCH1 expression, instead of the average value, within the microtumor promotes the invasiveness of microtumors.

**Figure 5.**
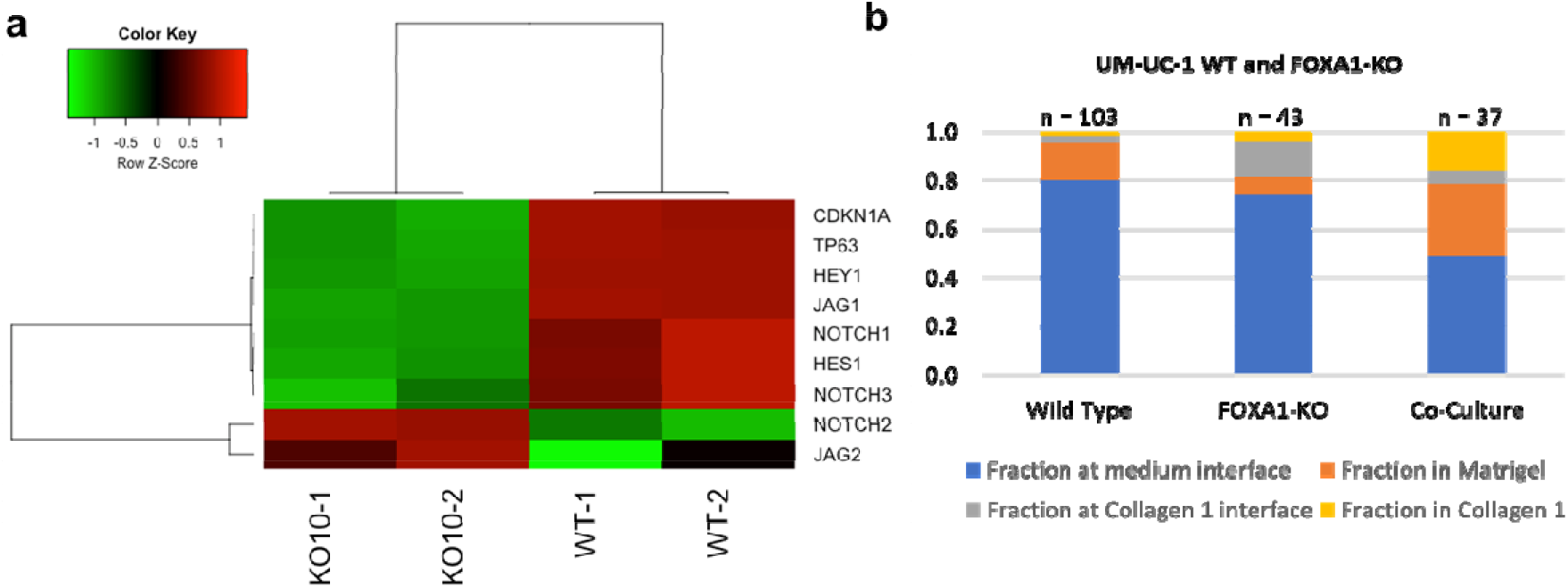
FOXA1 and NOTCH1 expression. (a) RNA-seq results indicate that NOTCH1 is downregulated in FOXA1 knockout UM-UC-1 cells. (b) Number of microtumors in each region for UM-UC-1 wild-type, FOXA1 knockout, and co-culture conditions.

### Computational modeling of mixed microtumor with variations in NOTCH1 expression

To evaluate how NOTCH1 variation may contribute to the enhanced invasiveness, we developed an agent-based computational model for evaluating the effects of NOTCH1 expression variations within the microtumor. The agent-based model considered the production, degradation, and cis-inhibition of NOTCH1 and DLL4 (Fortini, 2009). The production rate of NOTCH1 was modulated by the level of DLL4 of cells in contact, and the production rate of DLL4 was attenuated by the NOTCH1 activity (**Fig. 6a**). These interactions resulted in lateral inhibition of DLL4 expressing cells. When the maximum production rate of NOTCH1, *R_N_*, was uniform for all cells, lateral inhibition of NOTCH1-DLL4 created a pattern with regular spacing of DLL4 expressing cells surrounded by NOTCH1 expressing cells (**Fig. 6b**). The check box (or mosaic) pattern formed with various values of maximum NOTCH1 and DLL4 production rates (**Supplementary Fig. S5**). Lateral inhibition provides a robust mechanism for the formation of the checker box pattern, which is used to explain hair cell patterning during embryonic development (Shaya and Sprinzak, 2011).

**Figure 6.**
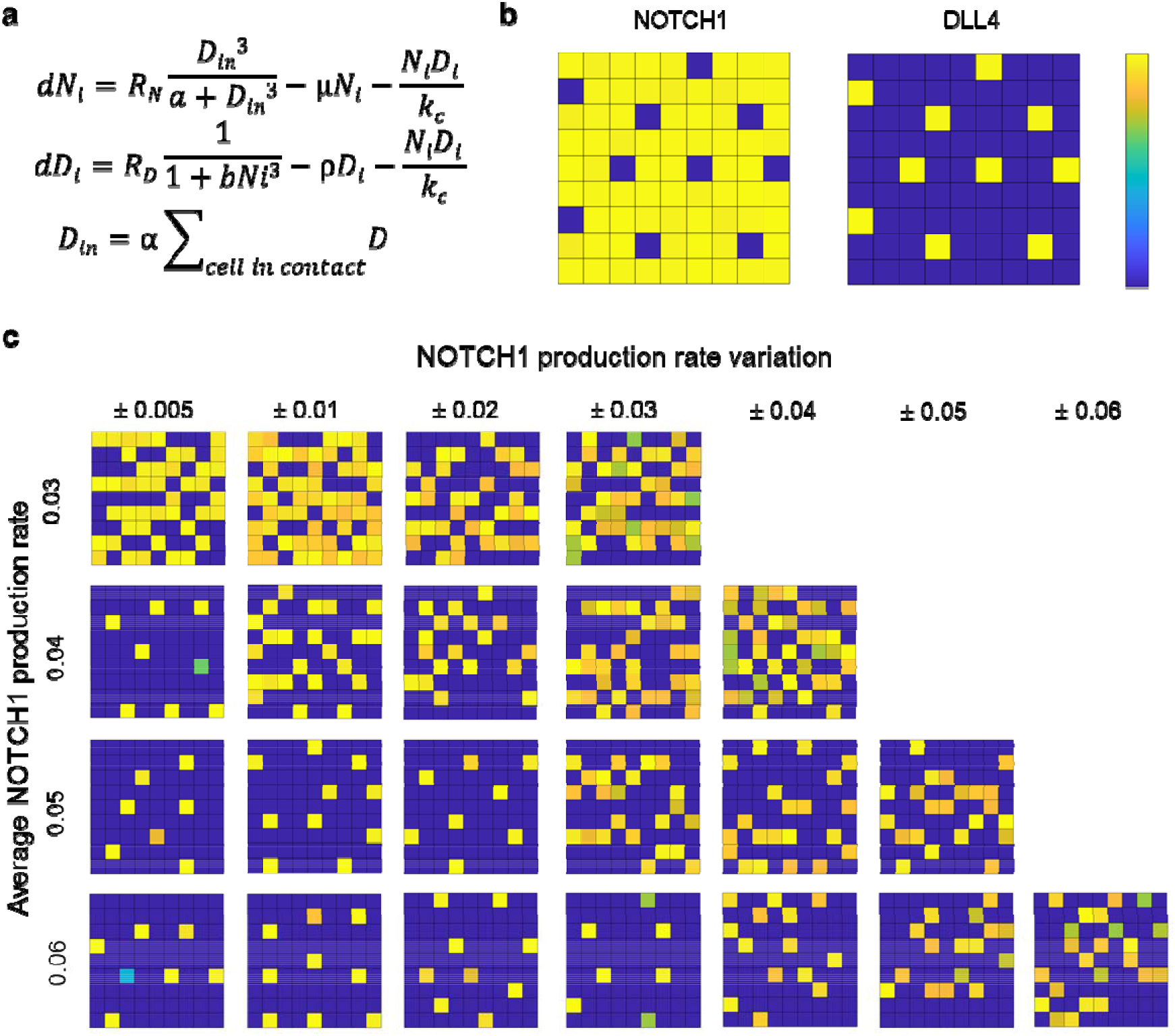
Agent-based computational modeling reveals NOTCH1 variation enhances DLL4 expression. (a) Differential equations employed in the agent-based computational model of NOTCH1-DLL4 expression. (b) Formation a checker box pattern of DLL4 expression with homogeneous NOTCH1 production. (c) Effects of increasing average NOTCH1 production rate and NOTCH1 variation on DLL4 expression within the microtumor. The grey scale indicates the expression of DLL4 in arbitrary units.

Using the agent-based model, we investigated the effect of NOTCH1 expression variations. By introducing some variation of NOTCH production rate, the checker box pattern was disrupted (**Supplementary Fig. S6**). Unlike the checker box pattern with DLL4 expressing cells surrounded by NOTCH1 expressing cells, clusters of DLL4 expressing cells and NOTCH1 expressing cells were formed and occupied in alternative regions. Examining the expression dynamics revealed that the cells committed into either NOTCH1 or DLL4 phenotypes with various equilibrium values (**Supplementary Fig. S6**). This observation suggests variation of NOTCH1 expression can have a significant effect on DLL4 activities. To understand the effect of variation NOTCH1 expression, we systematically adjusted the level of average NOTCH1 production rate and NOTCH1 production rate variation (**Fig. 6c**). With a high level of average NOTCH1 production rate, the checker box pattern was formed in most cases. Remarkably, a variation of the maximum NOTCH1 production rate was sufficient to modulate the checker box pattern to DLL4 clusters, despite that the average value was maintained constant. The level of variation of NOTCH1 expression increased with the cluster size. In addition, the NOTCH1 variation required for inducing the transition from checker box pattern to DLL4 clusters increased with the average NOTCH production rate (**Fig. 6c**). Even at a high average NOTCH1 production rate, a sufficiently large variation could disrupt the checker box pattern and created DLL4 clusters. The agent-based model suggests that variation of NOTCH1 expression can contribute to the formation of DLL4 expressing cell clusters with the microtumors. Importantly, the model provides a mechanistic explanation on how NOTCH1 variation, instead of average expression, promotes DLL4 expression within the mixed microtumor.

## Discussion

This study demonstrated a tumor bioengineering approach for investigating the influence of intratumoral cellular heterogeneity on cancer invasiveness. Self-assembling of microtumors allows systematic control the cell composition within the microtumors using established cell lines with luminal or basal transcriptional signatures. In addition, CRISPR/Cas9 gene editing and siRNA gene knockdown approaches enable us to test the role of subtype master regulators and molecular mechanisms important for invasion. The 3D invasion assay incorporates multiple layers of matrices representing bladder basement membrane and lamina propria. Matrigel and collagen matrices are commonly applied for mimicking the basement membrane and stromal ECM (Nguyen-Ngoc et al., 2012), and cancer cells can adopt various invasion modes depending on the microenvironment (Clark and Vignjevic, 2015; Friedl et al., 2012). Our invasion model resolves the potential of cancer cells to breach the basement membrane matrix and then subsequently invade the lamina propria. The ability to cross both layers suggests that the cancer subtype, or the combination of cancer subtypes, has the potential to progress to the muscularis propria, an important distinction in prognosis and management of bladder cancer. Unlike models that mixed multiple matrices (Anguiano et al., 2020; Carey et al., 2017), the multilayer approach revealed microtumor subpopulations with distinct abilities of invading different matrices, as indicated by microtumors trapped at the Matrigel-collagen interface. The tumor bioengineering approach provides a useful platform for characterizing the influence of intratumoral heterogeneity in a bladder mimicking environment. Logical continuations of this work would include further development of a multilayer tissue model to include other important components, such as fibroblasts, healthy urothelial cells, and microvasculature (or nutrient supply) and also incorporation of more cell lines and combinations in the multilayer model.

Using the tumor bioengineering approach, our results revealed intratumoral heterogeneity can enhance the invasiveness of microtumors. The heterogeneity mediated invasiveness was demonstrated using cell lines with distinct patterns of luminal (FOXA1, PPARG, and GATA3) and basal (KRT5 and KRT14) markers (**Supplementary Fig. S7**). For invasive cells (e.g., HT-1376), the invasion depth was not significantly enhanced suggesting that only some invasive cells in the microtumor are sufficient to induce collective invasion. This observation is in analogous to a heterotypic co-culture model of breast cancer, in which invasive malignant cells induce collective invasion of non-invasive cells (Carey et al., 2013). Conversely, the combination of SCaBER and UM-UC-1 was striking because of the significant enhancement of invasion observed in co-culture. Based on the invasive fraction, SCaBER and UM-UC-4 were actually the least invasive basal and luminal cell lines in this study. SCaBER and UM-UC-1 in co-culture showed much higher invasive potential than either cell line individually. When cultured separately, these cells exhibited only zero-to-low invasiveness in both single layer and multilayer models while combining the cell lines results in a large (> 40%) invasive fraction. The mixed microtumor displayed an enhanced capability of breaching through both Matrigel and collagen matrices.

Our results suggest that the heterogeneity induced invasiveness is associated with NOTCH1-DLL4 signaling. In particular, the invasiveness of microtumors were enhanced by the dissimilarity of NOTCH1 expression, instead of the absolute expression. The NOTCH expressions of SCaBER and UM-UC-1 cells were determined by the GNR-LNA single cell biosensor, which measured the expression of the microtumor in the 3D environment. NOTCH1 expression was significantly higher in UM-UC-1 microtumors than in SCaBER or co-culture, and mixed microtumor exhibited a large variation of NOTCH expression. Furthermore, transient knockdown of NOTCH1 in either UM-UC-1 and SCaBER, which promoted heterogeneity, resulted in a large invasive fraction (~35%) while NOTCH1 knockdown in both cells reduced the fraction of cells progressed to the collagen layer. The view that variation of NOTCH1 contributes to the enhanced invasiveness of heterogeneous is further supported by CRISPR-Cas9 gene knockout of FOXA-1. FOXA-1 KO reduced NOTCH1 and NOTCH1 target genes. While CRISPR KO of FOXA1 in UM-UC-1 did not have a strong effect on its invasiveness, co-culture of wild type and FOXA1 KO cells resulted in a clear increase in invasiveness. In particular, FOXA-1 KO itself did not lead to an enhanced in invasiveness compared to the wild type UM-UC-1 cells. The co-culture of UM-UC-1 and UM-UC-1 FOXA KO cells, which introduces a large variation in NOTCH expression within the microtumor, enhanced the invasiveness.

Multiple mechanisms may contribute to the intratumoral heterogeneity mediated tumor invasiveness. Our computational model revealed that variation in NOTCH1 can substantially enhance the expression of DLL4 and formed clusters of DLL4 cells. In the experiment, DLL4 knockdown in UM-UC-1 and SCaBER reduced the invasive fraction into the collagen layer for both individual culture and co-culture. The role of DLL4 in collective invasion is consistent with previous work examining NOTCH1-DLL4 signaling in HT-1376 bladder cancer cells and other cell models (Dean et al., 2016; Riahi et al., 2015; Torab et al., 2020). Nevertheless, DLL4 may not be a sufficient factor to induce the collective invasion process. Other ligands of NOTCH signaling, such as JAG1, may also be involved in the collective invasion process. For instance, epigenetic heterogeneity and JAG1 signaling were recently shown to jointly promote the persistence of filopodia via the MYO10 in invading cancer cells (Summerbell et al., 2020). The filopodia activity is necessary for creating migration tracks by micropatterning extracellular fibronectin in a 3D invasion model of Matrigel. Tumor heterogeneity, such as presence of luminal-basal and other molecular subtypes, may also enhance tumor invasiveness and survival capability via additional mechanisms. Future investigation will be required to clarify the functions of NOTCH signaling in collective invasion of cancer and examine other molecular processes associated with tumor heterogeneity.

## Methods

### Single layer invasion model

In this study, two 3D invasion models were used. A simplified tissue model representing the basement membrane matrix was used to screen bladder cancer cell lines for their ability to penetrate the basement membrane(Torab et al., 2020; Warrick et al., 2019). Briefly, a self-assembly process was used to generate a large number of bladder cancer microtumors on top of a layer of matrix representing the basement membrane composition. Growth factor reduced Matrigel (Corning) was diluted to 8 mg/mL in chilled, complete culture medium (Corning). Fluorescent beads (Spherotech) were then added at a volume ratio of 1:10,000 to label the Matrigel for imaging. 40 μL labeled Matrigel was applied to each well of a chilled glass-bottom 96-well plate (CellVis), which was then placed in a humidified cell incubator at 37°C, 5% CO_2_ for 30 min to solidify. Cells were then detached using 0.25% trypsin, 0.53 mM EDTA solution (Corning), suspended in complete culture medium at a concentration of 10^6^ cells/mL, and seeded atop the solidified Matrigel at a concentration of 10^4^ cells per well. The plate was then covered and returned to the cell culture incubator for 3 days before imaging.

### Multilayer invasion model

A multilayer tissue model representing the basement membrane (Matrigel) and lamina propria (Collagen 1) ECM layers was developed to study microtumor invasion for cell lines and combinations of cell lines that showed a high potential for invasion in the simplified model. Collagen type 1 from rat tail (SigmaAldrich) at a starting concentration of 4 mg/mL was diluted to 2 mg/mL in complete culture medium (Corning), and neutralized using 1M NaOH to a final pH of 7.75. The final concentration of fetal bovine serum (FBS, Corning) was 20%. 40 μL of collagen 1 was added to each well of a chilled glass-bottom 96-well plate (CellVis), which was then placed in a humidified cell incubator at 37°C, 5% CO_2_ for 60 min to solidify. Solidification was confirmed by visually inspecting for turbidity. Growth factor reduced Matrigel (Corning) was diluted to 5 mg/mL in chilled, complete culture medium (Corning). Fluorescent beads (Spherotech) were then added at a volume ratio of 1:10,000 to label the Matrigel for imaging. 10 μL labeled Matrigel was then carefully applied to each well on top of the solidified collagen 1 layer, and the entire 96-well plate was then returned to the cell incubator for 15 min to solidify. Cells were detached using 0.25% trypsin, 0.53 mM EDTA solution, and suspended in complete culture medium at a concentration of 10^6^ cells/mL, then seeded atop the solidified Matrigel layer at a concentration of 10^4^ cells per well.

### Cell culture

The bladder cancer cell lines RT4, UM-UC-3, SCaBER, HT-1197, HT-1376, and 5637 were obtained from ATCC, and UM-UC-1 was obtained from Sigma-Aldrich. RT4 cells were maintained in McCoy’s 5A culture medium and all other cell lines were maintained in MEM with 2 mM glutamine (Corning). Culture medium for HT-1197 and HT-1376 was supplemented with 1x non-essential amino acids and 1 mM sodium pyruvate (Gibco). All culture media were supplemented with 10% fetal bovine serum (Corning) and 1 μg/mL Gentamicin (Gibco). Cells were grown in 60 mm tissue culture dishes and were incubated at 37°C, 5% CO_2_ with 95% humidity. The cells were examined under a microscope on a daily basis, and the medium was renewed every 2 days. Cells were passaged at 70% confluence using 0.25% trypsin, 0.53 mM EDTA solution (Corning).

### Cell treatments

DLL4 siRNA, NOTCH1 siRNA, and non-targeting control siRNA were purchased from Santa Cruz Biotech. Transfection was performed in monolayer culture using Hiperfect transfection reagent (Qiagen) with an siRNA concentration of 30 pM for 24 hours prior to microtumor self-assembly. LNA fluorescent probes (Integrated DNA Technologies) targeting DLL4 and NOTCH1 were attached to MUTAB-coated GNRs (Nanopartz) in Tris-EDTA buffer to form GNR-LNA biosensor complex. Cells were incubated with the biosensor for 12 hours to allow for sufficient uptake. Cells were stained using CellTracker Green CMFDA or Red CMTPX (Invitrogen) at a concentration of 20 μM in PBS for 30 minutes. Stains were alternated in experimental replicates.

### Computational model

An agent-based computational model was developed for evaluating the effects of intratumoral variations of NOTCH1 expression. The numerical model was developed based on reported studies of NOTCH lateral inhibition(Cohen et al., 2010; Riahi et al., 2015). The model consisted of either 8 by 8 or 16 by 16 discretized elements (agents) to represent the microtumor and was solved in MATLAB or Octave. Parameters of the basic model were obtained from the previous study and were adjusted as indicated to evaluate the effects of NOTCH1 and DLL4 variations(Cohen et al., 2010). A periodic boundary condition was applied to the microtumor. Cells surrounding the first and second layers of a cell were considered in contact due to the dynamic filopodial activity of the cells(Cohen et al., 2010).

### CRISPR knockout

UM-UC-1 cells (2×10^5^) cells were transfected with 2.5 mg of HNF-3alpha CRISPR/Cas9 KO plasmid (Santa Cruz; sc-400743) using Lipofectamine 3000 (Thermo Fisher Scientific). After 48 hours, transfected cells were trypsinized and resuspended in PBS. Three GFP-positive cells were isolated via flow cytometry into single wells of 96-well plates (Corning) in 100 μl aliquots. Sorted cells were expanded and sequentially transferred to 24-well, 6-well dishes and T75 flasks (Corning). Finally, FOXA1 knockout was confirmed by qPCR and western blotting analysis.

### Statistics and data analysis

Invasion depth measurements (**Fig. 1f-m**) with unequal sample sizes were compared using a Kruskal-Wallis one-way ANOVA on ranks followed by a Tukey-Kramer post-hoc test. Biosensor intensity (**Fig. 3b-c**) measurements were compared using one-way ANOVA followed by Tukey’s HSD post-hoc test. Median invasion depth of co-culture microtumors was compared to that of each individual comprising cell line using a one-tailed Wilcoxon rank-sum test. To define a threshold for the invasive fraction, the K-means clustering algorithm (k=2) is applied. Statistical analysis was performed using the MATLAB statistics toolbox (Mathworks). For all figures, * indicates *p*<0.05, ** indicates *p*<0.01, and *** indicates *p*<0.001.

---

**Figure S1.**
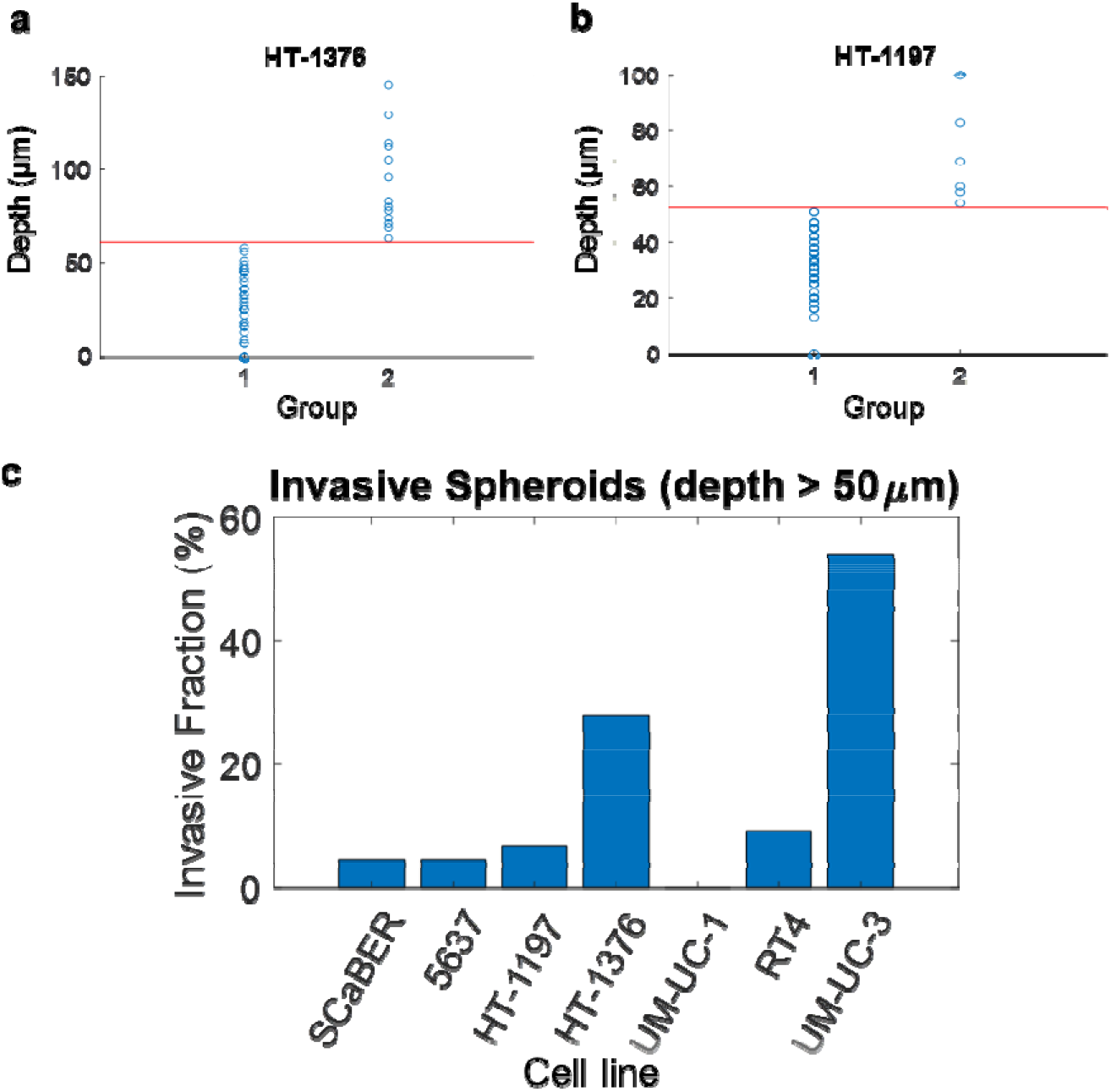
Clustering analysis defines the invasive fraction of microtumors in the single layer invasion model. (a-b) K-means clustering results for invasive microtumor depth in the single layer invasion model, 72 hours after seeding (k=2). In both cell line, the threshold value was determined to be approximately 50 μm. (C) Fraction of microtumors that invade Matrigel more than 50 μm. SCaBER and RT4 represent the least invasive cell lines in the basal and luminal subtypes, respectively. In addition to the luminal and basal cell lines, UM-UC-3, which is a highly invasive bladder cancer cell line and represents the neuroendocrine-like subtype, was included as a reference. Data are obtained from 3 independent experiments. n is at least 50 for each case.

**Figure S2.**
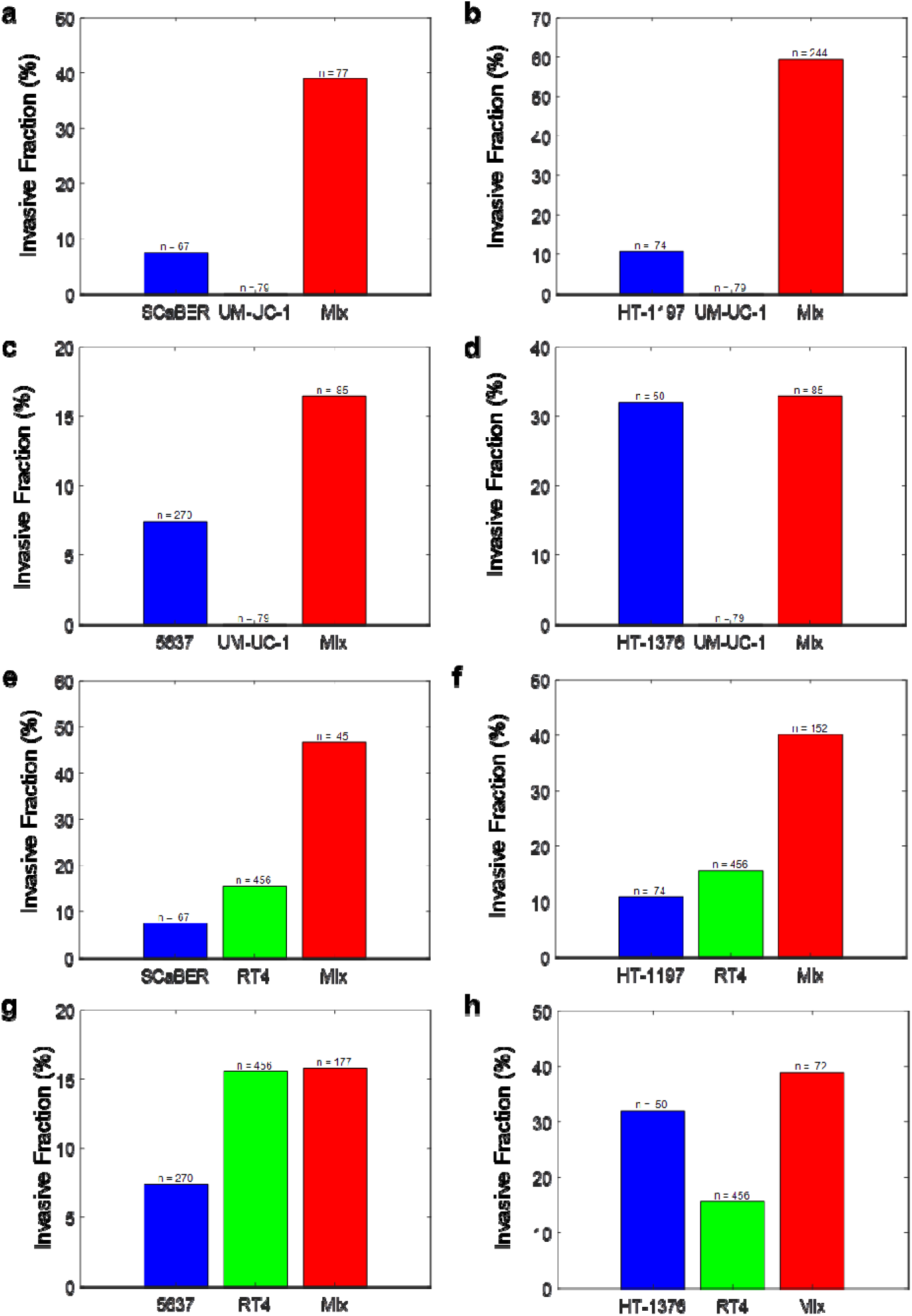
Co-culture enhances invasiveness of microtumors compared to individual cell lines. (a-h) Invasive fraction of mixed microtumors vs microtumors formed by individual cells lines. Data are obtained from 3 independent experiments. n is at least 45 for each case.

**Figure S3.**
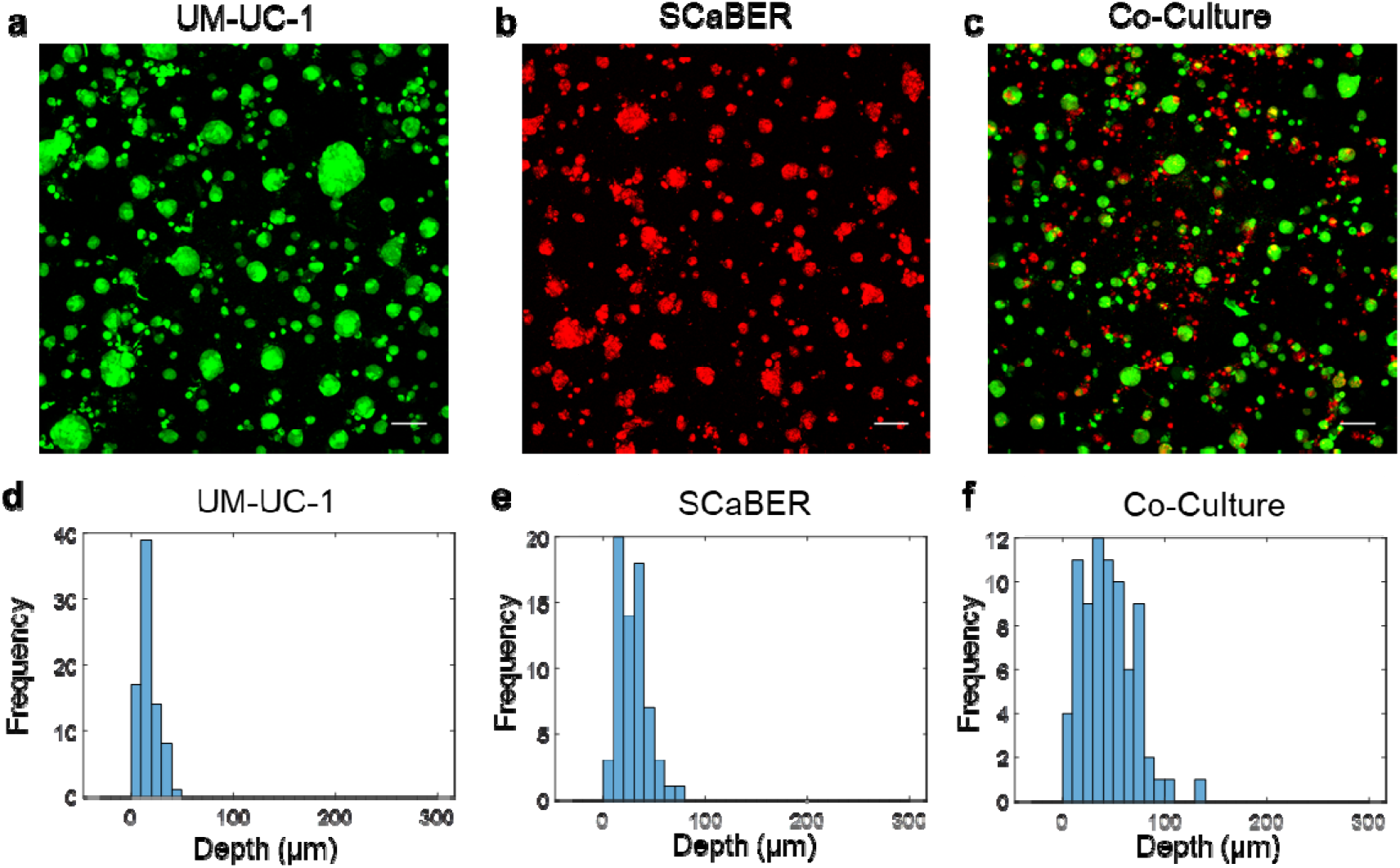
SCaBER and UM-UC-1 invasion of Matrigel. (a-c) Vertical projection views of the single layer invasion model. Scale bars, 100 μm. (d-e) Distribution of depth of microtumor invasion into Matrigel. Data are representative of 3 independent experiments.

**Figure S4.**
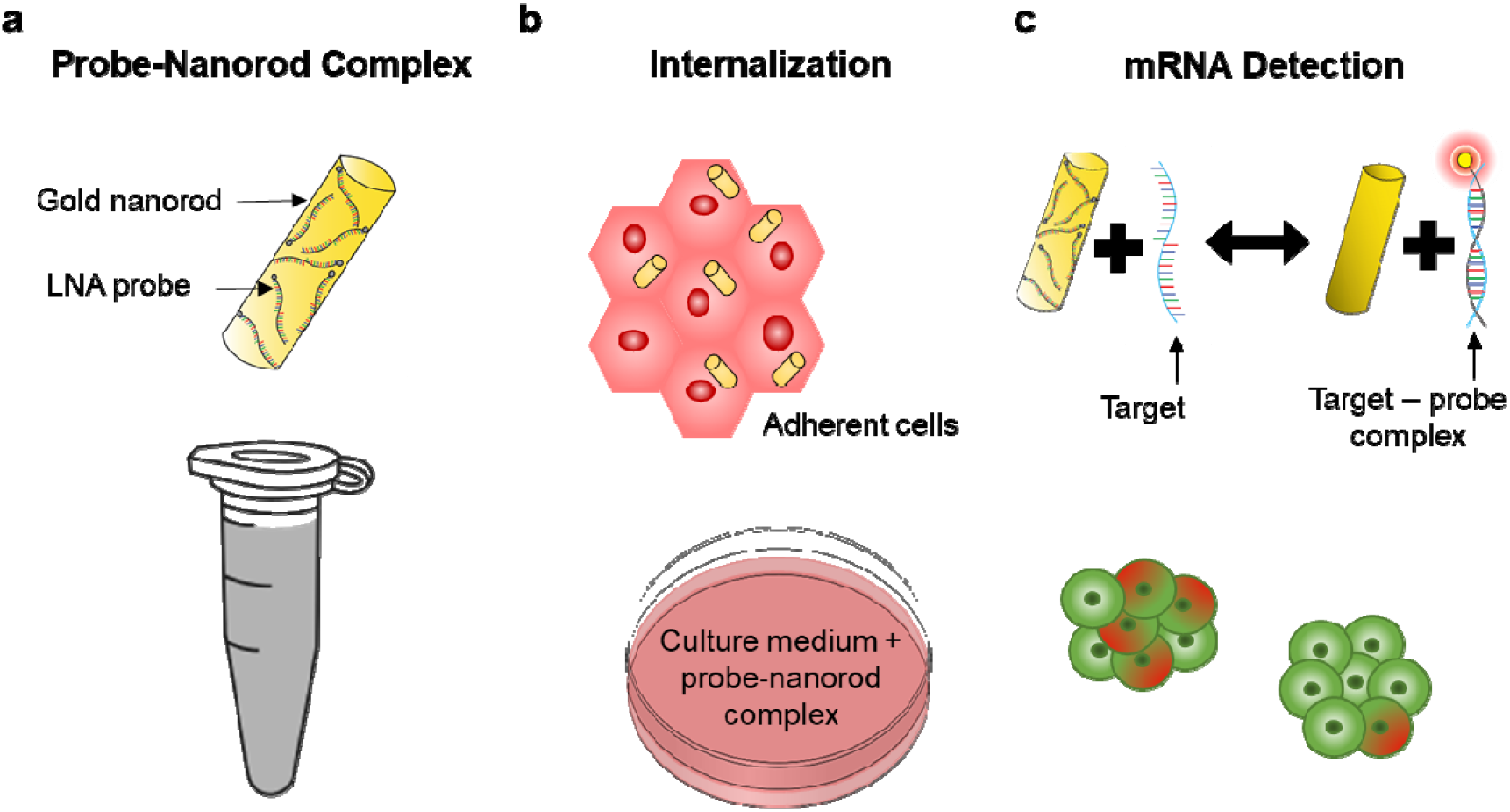
Live cell biosensors for measuring NOTCH1 and DLL4 expression in 3D microtumors. (a) Schematic of gold nanorod-locked nucleic acid (GNR-LNA) biosensors. (b) The biosensors are internalized into the cells to ensure uniform loading among the cells. (c) The cells are then self-assembled on ECM mimicking gel to form 3D microtumors.

**Figure S5.**
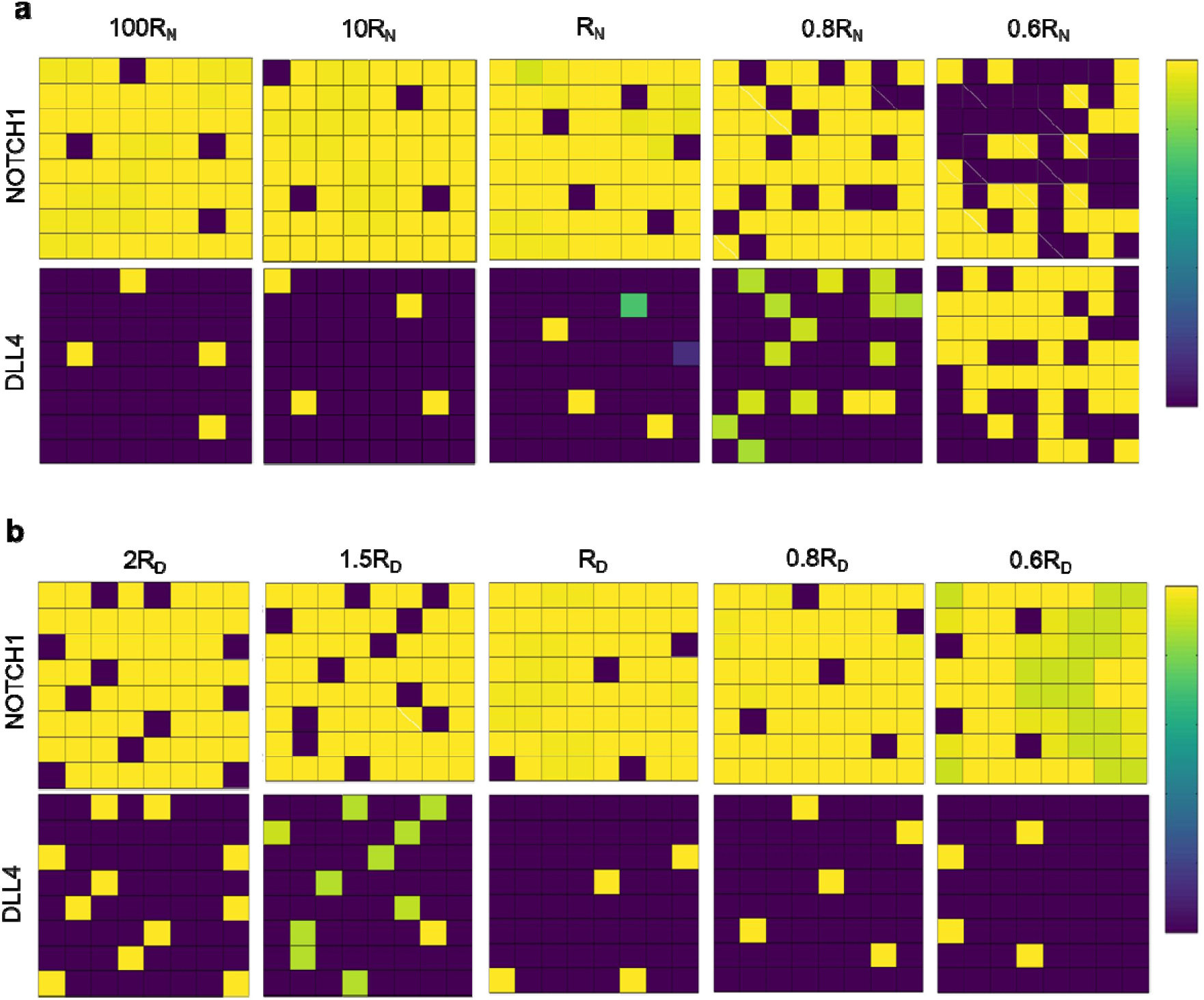
Computation modeling of NOTCH1 and DLL4. (a-b) Agent based model for studying the effects of NOTCH1 and DLL4 on pattern formation. Sensitivity analysis of (a) NOTCH1 production rates *R_N_* and (b) DLL4 production rate *R_D_* for evaluating the effects of NOTCH1 and DLL4 production rates on the spatial pattern. The base values of R_N_ and R_D_ were 0.01.

**Figure S6.**
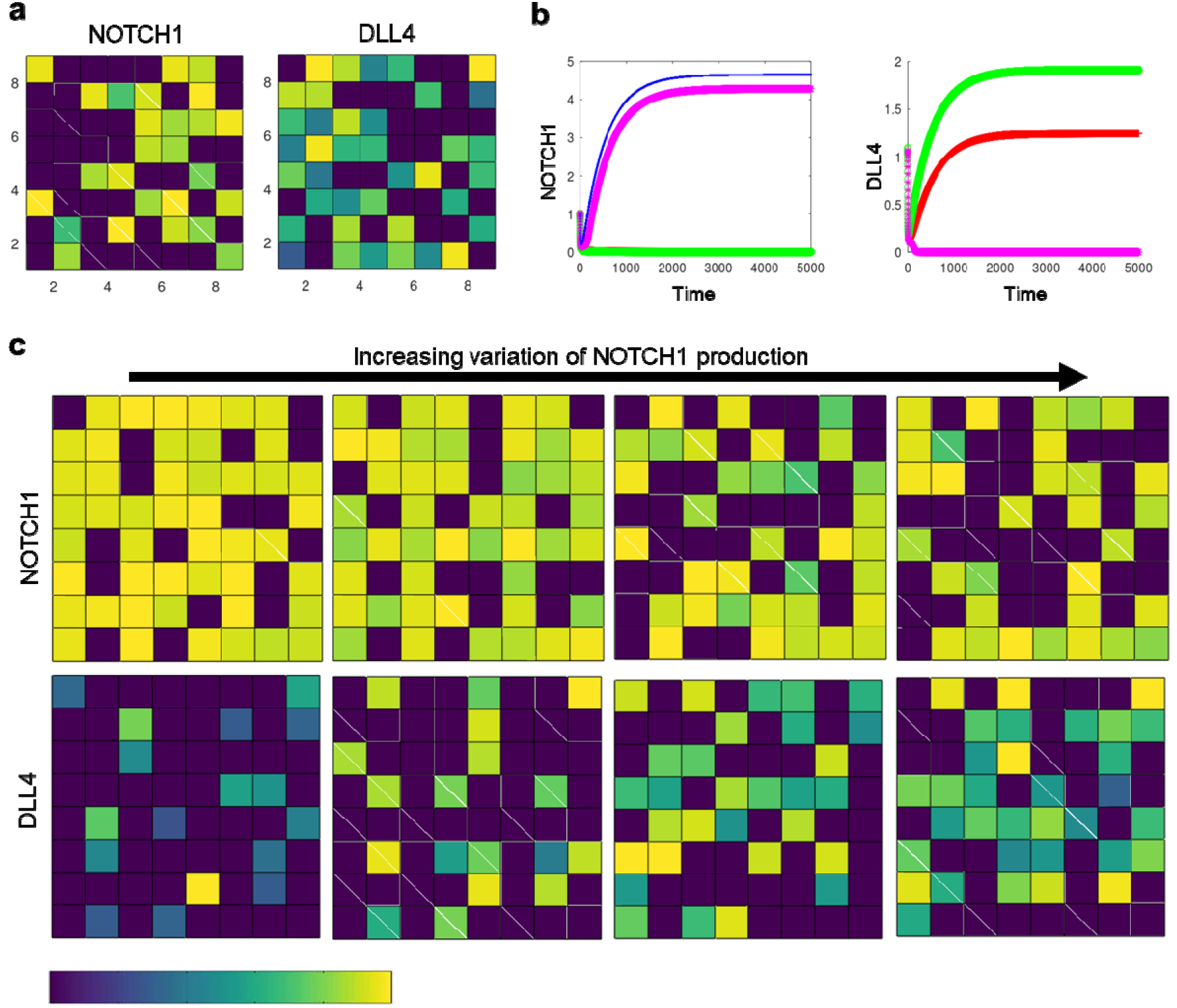
Computation modeling of DLL4 expression with variation of NOTCH1 production rate. (a) Introduction of NOTCH1 variation resulted in cluster of DLL4 expressing cells. (b) Tracking of NOTCH1 and DLL4 activities in representative cells committed to NOTCH1 or DLL4 phenotypes. (c) Effects of increasing NOTCH1 variation on DLL4 expression within the microtumor. The color bar shows the level of NOTCH1 and DLL4 in arbitrary units.

**Figure S7.**
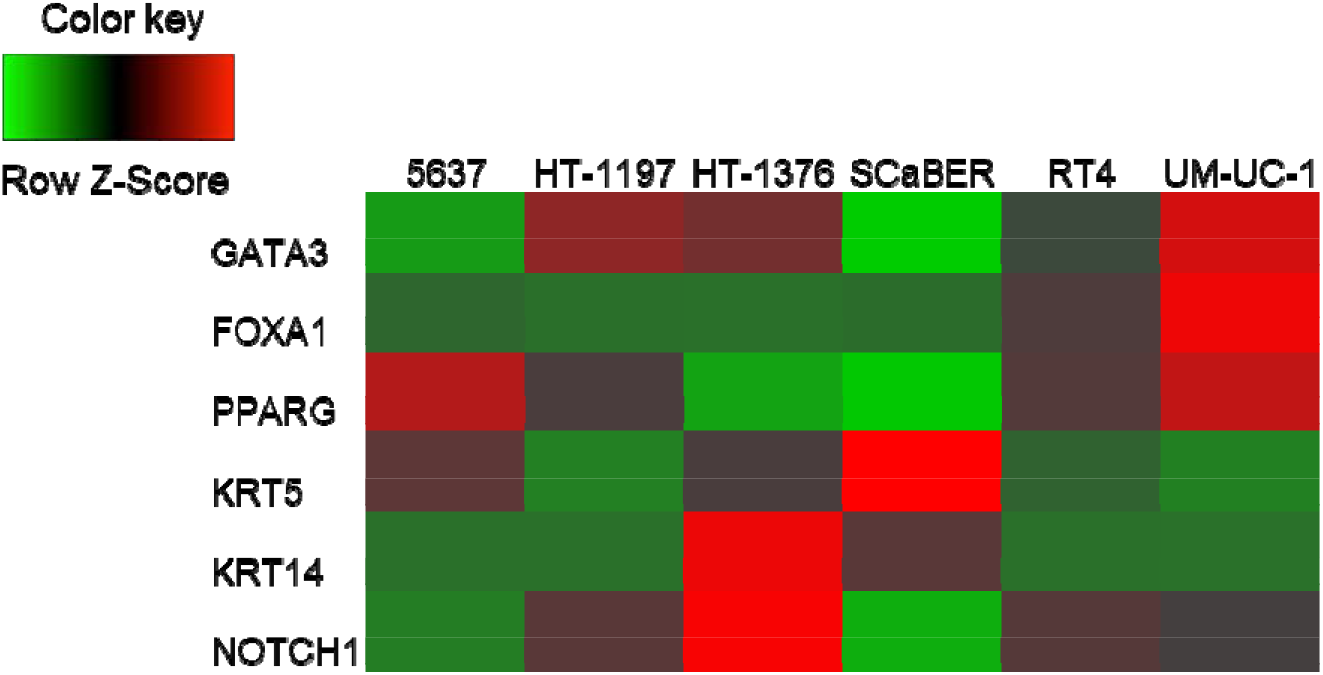
Comparison of cancer cell line encyclopedia RNA-seq data for luminal and basal cell lines. RT4 and UM-UC-1 express typical markers of luminal cells (GATA3, FOXA1, and PPARG) while 5673, HT-1197, HT-1376, SCaBER express typical markers of basal cells (KRT5 and KTR14). These cell lines, such as the UM-UC-1 and SCaBER pair, express distinct levels of NOTCH1 at the transcriptional level.

## Supplementary Movies

**Movie S1.** Heterogeneous microtumors in 3D microenvironment. Heterogeneous microtumors formed by bladder cancer cells of luminal papillary (UM-UC-1) and basal/squamous (SCaBER) subtypes. The Red: SCaBER; Green: UM-UC-1. 3D microtumors are approximately 50-100 μm in diameter.

**Movie S2.** Heterogeneous microtumors formed by bladder cancer cells of luminal papillary (UM-UC-1) and basal/squamous (SCaBER) subtypes. The Red: SCaBER; Green: UM-UC-1. 3D microtumors are approximately 50-100 μm in diameter.

**Movie S3.** Heterogeneous microtumors formed by bladder cancer cells of luminal papillary (UM-UC-1) and basal/squamous (SCaBER) subtypes. The Red: SCaBER; Green: UM-UC-1. 3D microtumors are approximately 50-100 μm in diameter.

## References

Anguiano, M., Morales, X., Castilla, C., Pena, A. R., Ederra, C., Martinez, M., Ariz, M., Esparza, M., Amaveda, H., Mora, M. et al. (2020). The use of mixed collagen-Matrigel matrices of increasing complexity recapitulates the biphasic role of cell adhesion in cancer cell migration: ECM sensing, remodeling and forces at the leading edge of cancer invasion. PLoS One 15, e0220019.

Cancer Genome Atlas Research, N. (2014). Comprehensive molecular characterization of urothelial bladder carcinoma. Nature 507, 315–22.

Carey, S. P., Martin, K. E. and Reinhart-King, C. A. (2017). Three-dimensional collagen matrix induces a mechanosensitive invasive epithelial phenotype. Sci Rep 7, 42088.

Carey, S. P., Starchenko, A., McGregor, A. L. and Reinhart-King, C. A. (2013). Leading malignant cells initiate collective epithelial cell invasion in a three-dimensional heterotypic tumor spheroid model. Clin Exp Metastasis 30, 615–30.

Cheung, K. J., Gabrielson, E., Werb, Z. and Ewald, A. J. (2013). Collective invasion in breast cancer requires a conserved basal epithelial program. Cell 155, 1639–51.

Choi, W., Porten, S., Kim, S., Willis, D., Plimack, E. R., Hoffman-Censits, J., Roth, B., Cheng, T., Tran, M., Lee, I. L. et al. (2014). Identification of distinct basal and luminal subtypes of muscle-invasive bladder cancer with different sensitivities to frontline chemotherapy. Cancer Cell 25, 152–65.

Clark, A. G. and Vignjevic, D. M. (2015). Modes of cancer cell invasion and the role of the microenvironment. Curr Opin Cell Biol 36, 13–22.

Cohen, M., Georgiou, M., Stevenson, N. L., Miodownik, M. and Baum, B. (2010). Dynamic Filopodia Transmit Intermittent Delta-Notch Signaling to Drive Pattern Refinement during Lateral Inhibition. Developmental Cell 19, 78–89.

Damrauer, J. S., Hoadley, K. A., Chism, D. D., Fan, C., Tiganelli, C. J., Wobker, S. E., Yeh, J. J., Milowsky, M. I., Iyer, G., Parker, J. S. et al. (2014). Intrinsic subtypes of highgrade bladder cancer reflect the hallmarks of breast cancer biology. Proc Natl Acad Sci U S A 111, 3110–5.

Dean, Z. S., Elias, P., Jamilpour, N., Utzinger, U. and Wong, P. K. (2016). Probing 3D Collective Cancer Invasion Using Double-Stranded Locked Nucleic Acid Biosensors. Anal Chem 88, 8902–8907.

Fortini, M. E. (2009). Notch Signaling: The Core Pathway and Its Posttranslational Regulation. Developmental Cell 16, 633–647.

Friedl, P., Locker, J., Sahai, E. and Segall, J. E. (2012). Classifying collective cancer cell invasion. Nat Cell Biol 14, 777–83.

Fujiyama, C., Jones, A., Fuggle, S., Bicknell, R., Cranston, D. and Harris, A. L. (2001). Human bladder cancer invasion model using rat bladder in vitro and its use to test mechanisms and therapeutic inhibitors of invasion. British Journal of Cancer 84, 558–564.

Goriki, A., Seiler, R., Wyatt, A. W., Contreras-Sanz, A., Bhat, A., Matsubara, A., Hayashi, T. and Black, P. C. (2018). Unravelling disparate roles of NOTCH in bladder cancer. Nat Rev Urol 15, 345–357.

Hwang, P. Y., Brenot, A., King, A. C., Longmore, G. D. and George, S. C. (2019). Randomly Distributed K14(+) Breast Tumor Cells Polarize to the Leading Edge and Guide Collective Migration in Response to Chemical and Mechanical Environmental Cues. Cancer Res 79, 1899–1912.

Jager, W., Moskalev, I., Janssen, C., Hayashi, T., Gust, K. M., Awrey, S. and Black, P. C. (2014). Minimally Invasive Establishment of Murine Orthotopic Bladder Xenografts. Jove-Journal of Visualized Experiments.

Kamoun, A., de Reynies, A., Allory, Y., Sjodahl, G., Robertson, A. G., Seiler, R., Hoadley, K. A., Groeneveld, C. S., Al-Ahmadie, H., Choi, W. et al. (2020). A Consensus Molecular Classification of Muscle-invasive Bladder Cancer. Eur Urol 77, 420–433.

Liu, Y., Bui, M. M. and Xu, B. (2017). Urothelial Carcinoma With Squamous Differentiation Is Associated With High Tumor Stage and Pelvic Lymph-Node Metastasis. Cancer Control 24, 78–82.

Luo, Y., Zhu, Y. T., Ma, L. L., Pang, S. Y., Wei, L. J., Lei, C. Y., He, C. W. and Tan, W. L. (2016). Characteristics of bladder transitional cell carcinoma with E-cadherin and N-cadherin double-negative expression. Oncol Lett 12, 530–536.

Maraver, A., Fernandez-Marcos, P. J., Cash, T. P., Mendez-Pertuz, M., Duenas, M., Maietta, P., Martinelli, P., Munoz-Martin, M., Martinez-Fernandez, M., Canamero, M. et al. (2015). NOTCH pathway inactivation promotes bladder cancer progression. J Clin Invest 125, 824–30.

Mariathasan, S., Turley, S. J., Nickles, D., Castiglioni, A., Yuen, K., Wang, Y., Kadel, E. E., III, Koeppen, H., Astarita, J. L., Cubas, R. et al. (2018). TGFbeta attenuates tumour response to PD-L1 blockade by contributing to exclusion of T cells. Nature 554, 544–548.

McGranahan, N. and Swanton, C. (2015). Biological and therapeutic impact of intratumor heterogeneity in cancer evolution. Cancer Cell 27, 15–26.

Nam, S., Lee, J., Brownfield, D. G. and Chaudhuri, O. (2016). Viscoplasticity Enables Mechanical Remodeling of Matrix by Cells. Biophys J 111, 2296–2308.

Nguyen-Ngoc, K. V., Cheung, K. J., Brenot, A., Shamir, E. R., Gray, R. S., Hines, W. C., Yaswen, P., Werb, Z. and Ewald, A. J. (2012). ECM microenvironment regulates collective migration and local dissemination in normal and malignant mammary epithelium. Proc Natl Acad Sci U S A 109, E2595–604.

Padmanaban, V., Tsehay, Y., Cheung, K. J., Ewald, A. J. and Bader, J. S. (2020). Between-tumor and within-tumor heterogeneity in invasive potential. PLoS Comput Biol 16, e1007464.

Qiu, M., Bao, W., Wang, J., Yang, T., He, X., Liao, Y. and Wan, X. (2014). FOXA1 promotes tumor cell proliferation through AR involving the Notch pathway in endometrial cancer. Bmc Cancer 14, 78.

Ramakrishnan, S., Huss, W., Foster, B., Ohm, J., Wang, J., Azabdaftari, G., Eng, K. H. and Woloszynska-Read, A. (2018). Transcriptional changes associated with in vivo growth of muscle-invasive bladder cancer cell lines in nude mice. Am J Clin Exp Urol 6, 138–148.

Rampias, T., Vgenopoulou, P., Avgeris, M., Polyzos, A., Stravodimos, K., Valavanis, C., Scorilas, A. and Klinakis, A. (2014). A new tumor suppressor role for the Notch pathway in bladder cancer. Nat Med 20, 1199–205.

Riahi, R., Sun, J., Wang, S., Long, M., Zhang, D. D. and Wong, P. K. (2015). Notch1-Dll4 signalling and mechanical force regulate leader cell formation during collective cell migration. Nat Commun 6, 6556.

Shaya, O. and Sprinzak, D. (2011). From Notch signaling to fine-grained patterning: Modeling meets experiments. Current Opinion in Genetics & Development 21, 732–739.

Siegel, R. L., Miller, K. D., Fuchs, H. E. and Jemal, A. (2021). Cancer Statistics, 2021. CA Cancer J Clin 71, 7–33.

Style, R. W., Hyland, C., Boltyanskiy, R., Wettlaufer, J. S. and Dufresne, E. R. (2013). Surface tension and contact with soft elastic solids. Nat Commun 4, 2728.

Summerbell, E. R., Mouw, J. K., Bell, J. S. K., Knippler, C. M., Pedro, B., Arnst, J. L., Khatib, T. O., Commander, R., Barwick, B. G., Konen, J. et al. (2020). Epigenetically heterogeneous tumor cells direct collective invasion through filopodia-driven fibronectin micropatterning. Science Advances 6, eaaz6197.

Tao, S., Wang, S., Moghaddam, S. J., Ooi, A., Chapman, E., Wong, P. K. and Zhang, D. D. (2014). Oncogenic KRAS confers chemoresistance by upregulating NRF2. Cancer Res 74, 7430–41.

Tien, J., Ghani, U., Dance, Y. W., Seibel, A. J., Karakan, M. C., Ekinci, K. L. and Nelson, C. M. (2020). Matrix Pore Size Governs Escape of Human Breast Cancer Cells from a Microtumor to an Empty Cavity. iScience 23, 101673.

Torab, P., Yan, Y., Yamashita, H., Warrick, J. I., Raman, J. D., DeGraff, D. J. and Wong, P. K. (2020). Three-Dimensional Microtumors for Probing Heterogeneity of Invasive Bladder Cancer. Anal Chem 92, 8768–8775.

Wang, S., Riahi, R., Li, N., Zhang, D. D. and Wong, P. K. (2015). Single Cell Nanobiosensors for Dynamic Gene Expression Profiling in Native Tissue Microenvironments. Adv Mater 27, 6034–8.

Warrick, J. I., Kaag, M., Raman, J. D., Chan, W., Tran, T., Kunchala, S., Shuman, L., DeGraff, D. and Chen, G. (2017). FOXA1 and CK14 as markers of luminal and basal subtypes in histologic variants of bladder cancer and their associated conventional urothelial carcinoma. Virchows Arch 471, 337–345.

Warrick, J. I., Sjodahl, G., Kaag, M., Raman, J. D., Merrill, S., Shuman, L., Chen, G., Walter, V. and DeGraff, D. J. (2019). Intratumoral Heterogeneity of Bladder Cancer by Molecular Subtypes and Histologic Variants. Eur Urol 75, 18–22.

Warrick, J. I., Walter, V., Yamashita, H., Shuman, L., Amponsa, V. O., Zheng, Z., Chan, W., Whitcomb, T. L., Yue, F., Iyyanki, T. et al. (2016). FOXA1, GATA3 and PPAR□Cooperate to Drive Luminal Subtype in Bladder Cancer: A Molecular Analysis of Established Human Cell Lines. Sci Rep 6, 38531.

Wisdom, K. M., Adebowale, K., Chang, J., Lee, J. Y., Nam, S., Desai, R., Rossen, N. S., Rafat, M., West, R. B., Hodgson, L. et al. (2018). Matrix mechanical plasticity regulates cancer cell migration through confining microenvironments. Nat Commun 9, 4144.

Yamashita, H., Kawasawa, Y. I., Shuman, L., Zheng, Z., Tran, T., Walter, V., Warrick, J. I., Chen, G., Al-Ahmadie, H., Kaag, M. et al. (2019). Repression of transcription factor AP-2 alpha by PPARgamma reveals a novel transcriptional circuit in basal-squamous bladder cancer. Oncogenesis 8, 69.

Zaman, M. H., Trapani, L. M., Sieminski, A. L., Mackellar, D., Gong, H., Kamm, R. D., Wells, A., Lauffenburger, D. A. and Matsudaira, P. (2006). Migration of tumor cells in 3D matrices is governed by matrix stiffness along with cell-matrix adhesion and proteolysis. Proc Natl Acad Sci U S A 103, 10889–94.

Zuiverloon, T. C. M., de Jong, F. C., Costello, J. C. and Theodorescu, D. (2018). Systematic Review: Characteristics and Preclinical Uses of Bladder Cancer Cell Lines. Bladder Cancer 4, 169–183.

